# Proteoglycan IMPG2 shapes the interphotoreceptor matrix and modulates vision

**DOI:** 10.1101/859116

**Authors:** Ezequiel M Salido, Visvanathan Ramamurthy

## Abstract

The extracellular matrix surrounding the photoreceptor neurons, interphotoreceptor matrix (IPM) is comprised of two unique proteoglycans: IP<u>M p</u>roteo<u>g</u>lycan 1 and 2 (IMPG1 and IMPG2). Although the functions of the IPM are not understood, patients with mutations in IMPG1/2 develop visual deficits with subretinal material accumulation. Here, we generated mouse models lacking IMPG1/2 to decipher the role of these proteoglycans and the pathological mechanisms that lead to vision loss. IMPG1 and IMPG2 occupy specific locations in the outer retina, and both proteoglycans are fundamental for the constitution of the IPM system. Mice lacking IMPG2 show abnormal accumulation of IMPG1, and in later stages, develop subretinal lesions and reduced visual function. Interestingly, removal of IMPG1-2 showed normal retinal morphology and function, suggesting that the aberrant localization of IMPG1 causes the alterations observed in IMPG2 KO mice. In conclusion, our results demonstrate the role of IMPG2 in shaping the IPM, shed light on the potential mechanisms leading to subretinal lesions, and show that the secreted proteoglycans depend on the extracellular matrix environment to properly integrate into the matrix.

## Introduction

In the central nervous system, the extracellular matrix (ECM) form highly organized extracellular structures from molecules synthesized by neurons and glial cells. ECMs plays fundamental roles in cortical circuit development and plasticity (Berardi, Pizzorusso et al., 2004, Hensch, 2005). Deficits in ECM are associated with epilepsy, autism, schizophrenia, Alzheimer and blindness (Arranz, Perkins et al., 2014, Ishikawa, Sawada et al., 2015, Nicholson & Hrabetova, 2017, Sorg, Berretta et al., 2016, Wen, Binder et al., 2018, Xie, Kang et al., 2013). In the retina, the ECM surrounding the photoreceptors outer segments (OS) and inner segment (IS) is called the interphotoreceptor matrix (IPM) and extends from the Müller cell apical microvilli to the apical retinal pigment epithelium (RPE) microvilli (Ishikawa et al., 2015). Retina pathologies as macular dystrophies, subretinal drusenoid deposits (SDD) and age-related macular degeneration (AMD) are associated to deficits in the IPM and present lesions in the subretinal space (Alten & Eter, 2015, Boddu, Lee et al., 2014, Chowers, Tiosano et al., 2015, Hogg, 2014, Sivaprasad, Bird et al., 2016, Spaide, Ooto et al., 2018, Wilde, Lakshmanan et al., 2016).

The IPM constitutes the central transit zone for nutrients and metabolites between the RPE and photoreceptors (Ishikawa et al., 2015). In addition, it is thought that the IPM also participates in photoreceptor disk turn over, retinoid transport, calcium buffer, photoreceptor alignment, adhesion of the retina, and cell to cell communication, including growth factor presentation (Garlipp, Nowak et al., 2012, Ishikawa et al., 2015, Lazarus & Hageman, 1992, Strauss, 2005).

The IPM is composed of soluble and insoluble molecules. Soluble protein such as the interphotoreceptor retinoid-binding protein (IRBP) is involved in the trafficking of retinoids between photoreceptors and RPE, and is the most abundant IPM molecule (Pfeffer, Wiggert et al., 1983). The insoluble molecules are proteoglycans and glycosaminoglycans (GAG) such as chondroitin sulfate (ChS) and Hyaluronic acid (HA) (Clark, Keenan et al., 2011, Hauck, Schoeffmann et al., 2005, Hollyfield, Rayborn et al., 1998, Keenan, Clark et al., 2012, Tien, Rayborn et al., 1992). In comparison to other ECM, the IPM does not have collagen or large quantities of classical ChS proteoglycans such as aggrecan and neurocan (Ishikawa et al., 2015). Instead, the IPM has two particular ChS proteoglycans, the interphotoreceptor matrix proteoglycan 1 and 2 (IMPG1 and IMPG2). IMPG1 also referred as SPACR (sialoprotein associated with cones and rods) or IPM150 (Hollyfield, Rayborn et al., 2001, Kuehn & Hageman, 1999a, Lee, Chen et al., 2000) and IMPG2, also referred as SPACRCAN (sialoprotein associated with cones and rods proteoglycan) or IPM200 (Acharya, Foletta et al., 2000, Chen, Lee et al., 2003, Hollyfield et al., 2001, Kuehn & Hageman, 1999b). Both proteins are synthesized in the photoreceptors and their expression levels correlate with the maturation of photoreceptor OS (Felemban, Dorgau et al., 2018, Foletta, Nishiyama et al., 2001). The proteoglycans IMPG1 and IMPG2 are structurally similar. IMPG2 is larger than IMPG1 with a putative transmembrane domain that may aid in the attachment of IMPG2 to the cell membrane (Fig. 1). Both IMPG molecules have ChS attached and predicted attachment sites for HA (Acharya et al., 2000, Chen et al., 2003, Hollyfield, Rayborn et al., 1999).

**Figure 1.**
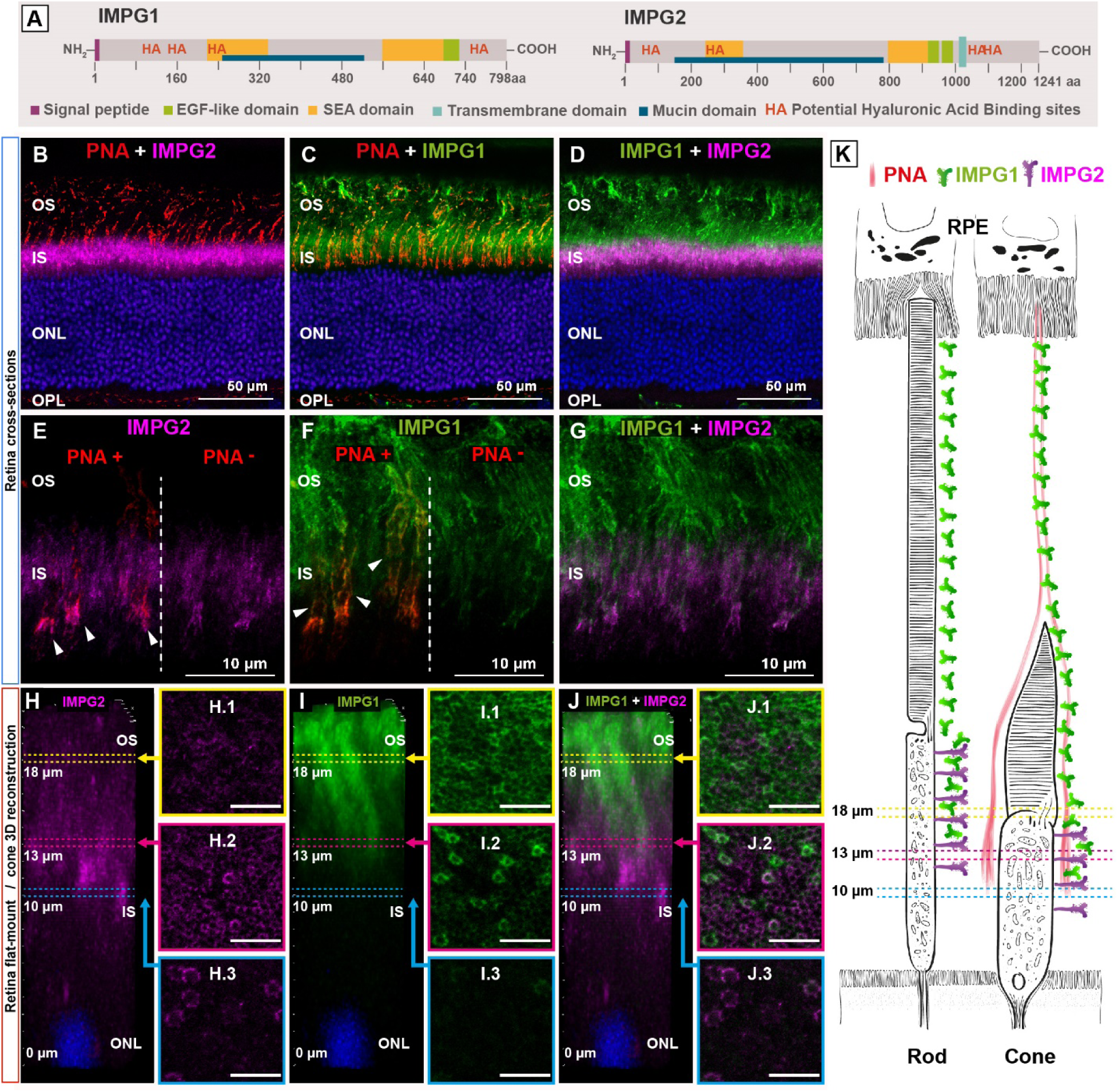
Localization of IMPG1 and IMPG2 proteoglycans in the IPM. Immunohistochemistry using retina from wild-type mice with IMPG1 (green) and IMPG2 (magenta) antibodies and cone-specific lectin, Peanut Agglutinin (PNA) (red). (A) Predicted structural domains of IMPG1 and IMPG2 proteins. Both IMPG1 and IMPG2 proteins have abundant O and N-glycosylation sites as well as potential hyaluronic acid attachment sites. SEA: sperm protein, enterokinase, agrin. EGF: epidermal growth factor. HA: hyaluronic acid (B-D) Retinal cross-sections from 45-day old animals imaged at low magnification. (E-G) Retinal cross-sections imaged at higher magnification. Arrowheads indicate Co-localization of PNA with IMPG2 and IMPG1 in panel E and F respectively. (H-J) Retina flat-mount 3D reconstruction focused on a single cone. On the right side of the 3D reconstruction image, flat mount raw confocal images used for 3D reconstruction selected at three different heights from the Outer Nuclear Layer (ONL) (10, 13 and 18 µm). There is a robust cone-IMPG2 stain at 10 µm above the ONL (H.3), and a weak IMPG1 mark (I.3). At 13 µm, rods and cones stains with IMPG2 and IMPG1 (H.2, I.2, J.2). The figure J.2 light up the cone-IMPG1 mark in green, surrounded by a smaller reticulated magenta pattern that belongs to rod-IMPG2 staining. At 18 µm height, IMPG2 stain is faint and unstructured (H.1), while IMPG1 mark is strong with a uniform reticular pattern (I.1) (Scale bars H-J, 10 μm). (K) Schematic representation of IMPG1, IMPG2, and PNA in cones and rods, combined with a height reference of the sections showed in subfigures H, I, and J. Pictures are representative of three independent experiments. OS: Outer Segment; IS: Inner Segment.

Humans with mutations in IMPG1 or IMPG2 can develop vision loss characterized by the accumulation of material in the macula subretinal space, a clinic symptom related with age foveomacular vitelliform dystrophy (AFVD), SDD and AMD (Bandah-Rozenfeld, Collin et al., 2010, Boddu et al., 2014, Brandl, Schulz et al., 2017, Chowers et al., 2015, Guziewicz, Sinha et al., 2017, Meunier, Manes et al., 2014, van Huet, Collin et al., 2014, Wilde et al., 2016, Zweifel, Spaide et al., 2011). Despite the implied role of IMPG1 and IMPG2 proteoglycans for our vision, little is known about the function of these proteins or the pathophysiology of the disease caused by mutations in these IMPGs. In this work, we delved into the IPM physiology, focusing on the role of IMPG1 and IMPG2, as well as into the pathophysiological mechanisms underneath visual impairment caused by defects in these proteins. For this purpose, we developed three global knockouts (KO) mice lines, IMPG1^-/-^, IMPG2^-/-^, and IMPG1-2 double KO. Our results indicate that IMPG2 proteoglycan is located at the IS and is required for the localization of IMPG1 in the OS. In the absence of IMPG2, IMPG1 accumulated between the OS and RPE leading to subretinal lesions, microglial activation, and decrease in visual function.

## Results

### Localization of photoreceptor-specific IMPGs proteoglycans

IMPG1 and 2 are photoreceptor-specific proteins within the retina (Acharya et al., 2000, Kuehn & Hageman, 1999a). A closer inspection of the IMPG protein sequence shows the presence of the N-terminal signal sequence, suggesting that these proteins are exported from the photoreceptor cells (Fig. 1.A). To determine the location of IMPGs in the murine retina, we performed an IHC analysis (Fig. 1). We found distinct patterns of IMPG1 and 2 staining surrounding the photoreceptor cells: specifically, IMPG2 is located in the distal regions of the IS (Fig. 1 B), while IMPG1 is found surrounding the IS and OS (Fig. 1 C). A magnified image (Fig. 1 E, F, G) shows IMPG1 and IMPG2 surrounding rods and cones. The lectin Peanut Agglutinin (PNA), an established cone-specific IPM marker colocalizes with both IMPG1 and IMPG2 (Fig. 1 E, F arrows).

In order to highlight the differences in localization of IMPGs between rods and cones, we performed flat-mount transversal sections of the retina and 3D reconstruction focusing on single cones (Fig. 2 H, I, J and the corresponding sub-rectangles). The IMPG2 stain appears 10 µm distal from the outer nuclear layer (ONL) as rings surrounding the cones (H3). At 13 µm, IMPG2 is expressed as smaller rings around rods in between cones (H2), further up, at 18 µm, IMPG2 staining is diffuse (H1). In contrast, weak staining of IMPG1 rings can be found around cones at 10 µm above the ONL (I3), where it co-localizes with IMPG2 (J3). At 13 µm, robust staining for IMPG1 around cones and faint IMPG1 around rods was noted. Further up at the OS level, this pattern disappears, and the IMPG1 stain is consistently strong. The superimposition of IMPG1 and IMPG2 images (panels J) highlight the transition from IMPG2 to IMPG1 and the different planes between cone and rod IMPGs. In this sense, Sub-rectangle I2 shows the cone-IMPG1 in green passing through a magenta honeycomb shape of rod-IMPG2 (J2). Supplementary movie 1 shows a 3D reconstruction video of IMPGs localization in wild-type retinas that help to visualize the spatial relationships among IMPG proteoglycans and between rods and cones. In summary, our results demonstrate that IMPG1 and 2 have distinct pattern distribution in the IPM and both, cone and rods, are surrounded by IMPG proteoglycans.

**Figure 2.**
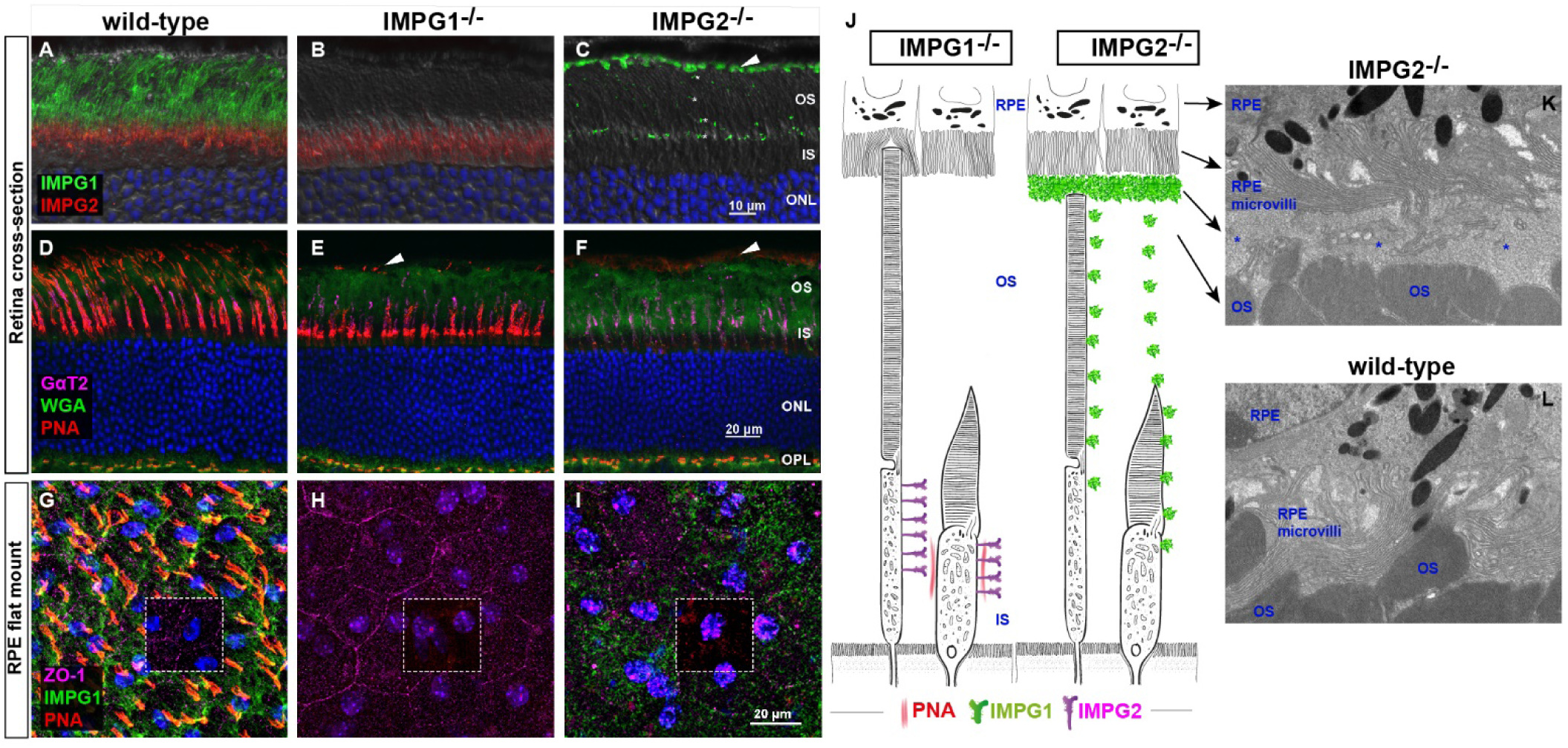
Mislocalization of IMPG1 in the absence of IMPG2. (A-I) Immunohistochemistry of the proteoglycans IMPG1, IMPG2 and PNA lectin on wild-type (left panel), IMPG1 (middle panel) and IMPG2 KO (right panel) mice. (B) IMPG1 KO mice show an absence of IMPG1 (green) and preserved IMPG2 (red) stain at the inner segment (IS). (C) Lack of IMPG2 staining in IMPG2 KO mice and accumulation of IMPG1 scattered throughout the outer segment (OS) (asterisks) and at the boundary between the RPE and the OS (arrow). (E) The disappearance of cone-specific lectin, PNA (red) at the OS of IMPG1 KO mice, leaving a small faint PNA mark at the RPE-OS boundary (arrow). In contrast, the lectin Wheat Germ Agglutinin (WGA) (green), and the cell cone marker, cone transducin protein, Gαt2, (magenta) are normal. (F) In IMPG2 KO retinas, WGA and Gαt2 staining extend normally, but PNA stain is absent or faint while some mislocalized PNA stains at the boundary between RPE and OS (arrow). PNA staining in unaltered at the outer plexiform layer (OPL) in all groups. (G-I) RPE flat-mount stained against PNA (red), IMPG1(green), and ZO-1 (magenta). (G) wild-type mice have a structured IMPG1 and PNA above RPE cells. (H) IMPG1 KO lacks IMPG1 and shows a very faint PNA stain above RPE. (I) IMPG2 KO shows a dense, unorganized IMPG1 staining with faint PNA above the RPE cells. Pictures are representative of three independent experiments. All images were obtained at 60X magnification using 45-days-old mice. (J) Schematic representation of the finding obtained from IMPG 1 and 2 KO mice. (K-L) Electron microscopy images centered in the intersection between photoreceptors outer segments (OS) and RPE microvilli. (K) IMPG2 KO retinas show a separation between OS and RPE filled by an unstructured material (asterisks). (L) Wild-type retinas exhibit close contact between OS and RPE microvilli. Data is representative of three animals at 5-months-old from 3 different littermates.

### IMPG1 location depends on IMPG2

To study the role of IMPG1 and IMPG2 in retinal physiology, we generated knockout (KO) mouse models using CRISPR-Cas9 technology. In IMPG1 mutant, Cas9 nuclease excision leads to a 10-nucleotide deletion in exon 2 producing a premature stop codon in exon 3. Similarly, in IMPG2 mutant, a 7-nucleotide deletion in exon 3 leads to premature termination on the same exon. In both cases, we expected non-sense mediated decay and consequent loss of IMPG1 and 2 proteins (*SI Appendix* Fig. S1). The IHC analysis confirmed the absence of IMPG1 or IMPG2 proteoglycans in IMPG1 and IMPG2 KO mice, respectively. These studies demonstrate the specificity of the antibodies used, and validate the knockouts generated by CRISPR-Cas9 methodology (Fig. 2A, B, C).

We observed that the distribution of IMPG2 at the IS was unaltered in the absence of IMPG1 (Fig. 2B). Interestingly, in IMPG2 KO mice, IMPG1 accumulates at the boundary between the OS and RPE, in addition to a scattered staining pattern through the OS (Fig. 2C). In addition, the cone-specific marker PNA disappeared around the OS in the absence of IMPG1 but remained at the IS region mimicking the IMPG2 staining pattern (Fig. 2E, B). In IMPG2 KO retina, PNA was mislocalized at the RPE-OS boundary like the aberrantly located IMPG1 (Fig. 2F, C). A 3D video reconstruction of flat-mounted retina from wild-type and mutant mice illustrate the absence of PNA in the OS of IMPG1 KO and the sporadic presence of IMPG1 in IMPG2 KO mice (Supplementary movie 2).

To further study the mislocalized IMPG1 and PNA at the boundary between the OS and the RPE, we performed an IHC analysis on flat-mount RPE cells. Wild-type tissue shows an organized and regularly spaced distribution of IMPG1 and PNA in contact with the RPE cells (Fig. 2G). However, flat mount from IMPG1 KO mice shows the absence of IMPG1 and a diffuse pattern of PNA staining (Fig. 2H). IMPG2 KO mice reveal a dense unstructured IMPG1 and PNA staining on the RPE surface (Fig. 2I). These results show a crucial role of IMPG2 proteoglycan in the IMPG1 distribution across the retina and the direct relationship between IMPG proteoglycans and PNA lectin staining.

### Subretinal material accumulation in IMPG2 KO mice

Next, we speculated that the misaccumulation of IMPG1 proteoglycan in IMPG2 KO retina affects the interaction between RPE microvilli and the photoreceptor OS. A magnified view by electron microscopy shows the accumulation of heterogeneous material that separates the RPE microvilli from the OS in IMPG2 KO mice (Fig. 2K and *SI Appendix* Fig. S2). In contrast, wild-type animals show the RPE microvilli in intimate contact with the OS (Fig. 2L). The IMPG1 KO retinas did not show any detectable changes in the RPE-OS interface.

### IMPG2 KO mice develop subretinal lesions

Patients with mutations linked to IMPGs proteoglycans develop visual impairment with punctate subretinal macular bodies detected by spectral domain optical coherence tomography (SD-OCT). Therefore, we investigated the retinal morphology using SD-OCT imaging in five-month-old IMPG2 KO mice. We found that most animals develop one or two hyperreflective clumps in the subretinal space (Fig. 3). These lesions above the RPE resemble the human lesions described as subretinal vitelliform lesions (Fig 3 A and A1, A3 arrows) (Meunier et al., 2014). The histological analysis of the SD-OCT studied retina confirmed these findings and showed a prominent bulge of fibrous materials separating the retina from the RPE and cells infiltrating in the subretinal space (Fig 3B, C, D). Moreover, the histological serial sections enabled the identification of several smaller subretinal material accumulation in IMPG2 KO retina that were difficult to detect using *in vivo* SD-OCT technique (Fig. 3A.2 and C). In contrast to IMPG2 KO retinas, no detectable alterations or subretinal accumulation were observed by OCT or histological analyses in littermate wild-type controls or IMPG1 KO animals. These results show that IMPG2 KO mice develop lesions that resemble the subretinal lesions found in humans with IMPG mutations.

**Figure 3.**
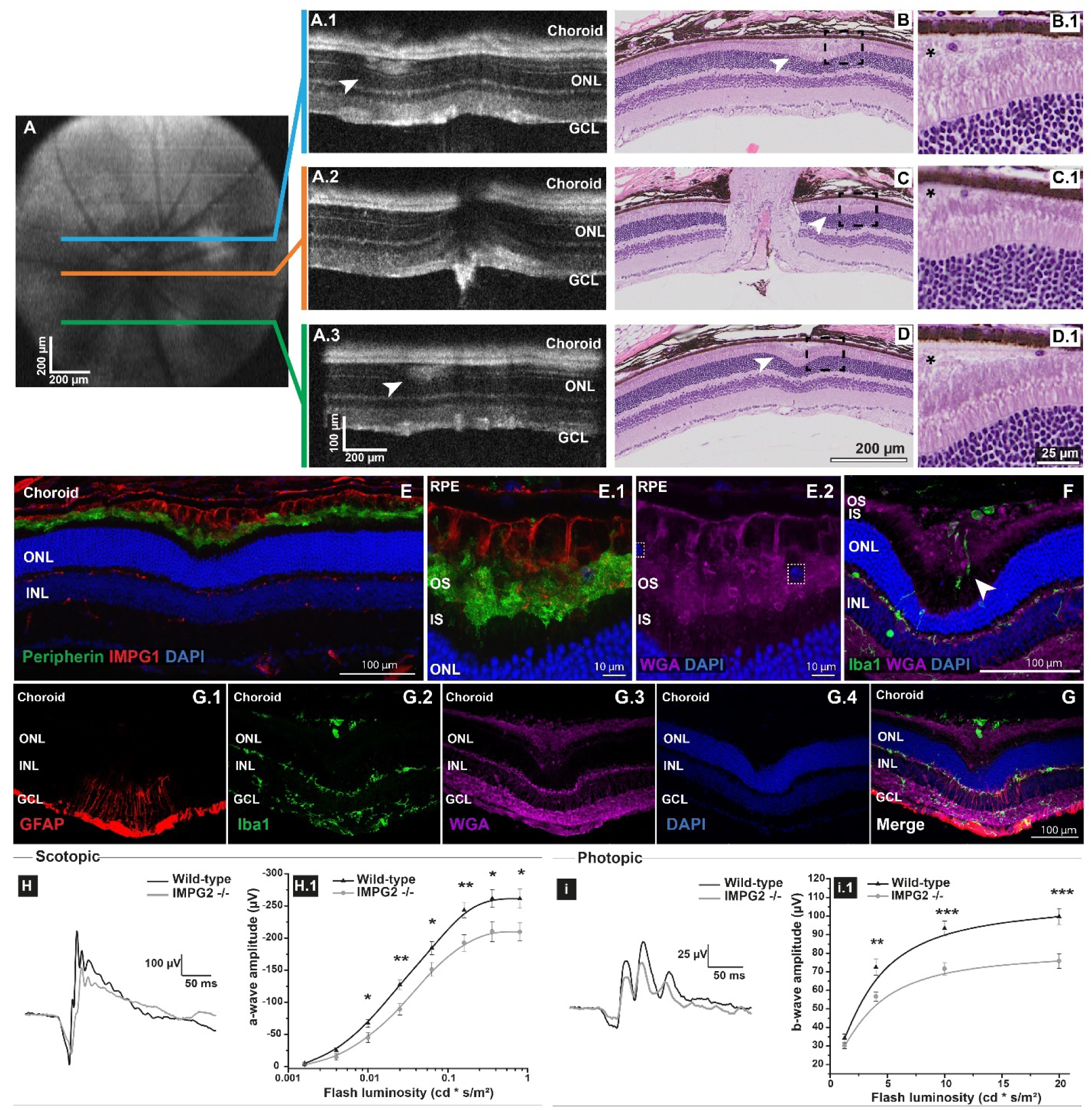
IMPG2 KO mice develop subretinal lesions. (A)Representative study of *in-vivo* spectral domain optical coherence tomography (SD-OCT) in IMPG2 KO. En-face image centered at the photoreceptors layer with color indicated orientations of cross-sectional B-scans shown in A.1, A.2 and A.3. (A.1, A.3) Subretinal hyperreflective nodules compromising the IS and OS layer, clinically referred as subretinal vitelliform lesions in humans (arrowheads). (B-D) H&E serial cross-sections of the same SD-OCT in Panel A. The subretinal lesions (arrowheads) exhibit a dome-shaped deformation on the retina. Magnification in the lesions shows an accumulation of eosinophil material between the OS and the RPE with the presence of infiltered cells (asterisks at B.1, C.1, D.1). Data is representative of eleven IMPG2 KO mice with subretinal lesions detected at 5-months-old. (E) Birds eye view of IHC of a subretinal lesion stained against IMPG1 (red) and the OS marker, peripherin (green). (E.1, E.2) Magnified view of the subretinal lesion shows IMPG1 accumulation over the OS and infiltrated cells stained with DAPI (white box) and WGA lectin (magenta). (F) Microglia infiltration stained with Iba1 (green) antibody migrating to the outer nuclear layer (ONL), IS and OS (arrow) and the subretinal space. (G) GFAP (red) reactivity circumscribed to a subretinal lesion (G.1), panel G shows the merged image of GFAP, Iba1 (green), WGA (magenta), and DAPI. (H-I) Reduced electroretinography (ERG) response in 8-months-old IMPG2 KO mice. Scotopic (rods) and photopic (cones) ERG responses on wild-type (black line) and IMPG2 knockout (gray line) mice, representative ERG traces at 0.16 and 10 Cd.s/m2 flash intensity respective. (H.1) Scotopic a-wave sensitivity curve. (I.1) Photopic b-wave responses from light-adapted mice (30 Cd/m2) to four increasing flash intensity. Data points are mean ± SEM of N=16 eyes, two-way ANOVA (8 mice per group from 3 different littermates), P values indicated as: ∗ P < 0.05, ∗∗ P < 0.01, ∗∗∗ P < 0.001.

### Subretinal lesions are rich in IMPG1 proteoglycan and activated microglia

Next, we hypothesized that the subretinal lesions are due to excessive accumulation of IMPG1 proteoglycan. To study the composition of these lesions, we stained cross-sections of subretinal lesions with IMPG1 antibody. The lesions showed distinct staining for IMPG1 in between the RPE and the OS (Fig. 3E, E.1, *SI Appendix* Fig. S3). In addition, the lesions show cell nuclei stained with DAPI (Fig. 3E.2 rectangles). Subsequently, we explored whether the cell nuclei in the subretinal lesions could be due to retinal microglia migrating to the area. Microglia were detected using an antibody against microglia/macrophage-specific calcium-binding protein (Iba1). In healthy retinas, microglial cells (Iba1+) displayed a ramified morphology, and are located specifically in the ganglion cell, and inner and outer plexiform layer (*SI Appendix* Fig. S3), as previously reported (Okunuki, Mukai et al., 2018). Interestingly, in IMPG2 KO mice we observed Iba1-positive cells invading the photoreceptor layers and the subretinal lesions (Fig. 3F, 4G). Then, we investigated whether macroglia also reacts to the subretinal lesion by staining against glial fibrillary acidic protein (GFAP). Müller reactive gliosis was circumscribed to the retina proximate to the lesions (Fig. 3G).

**Figure 4.**
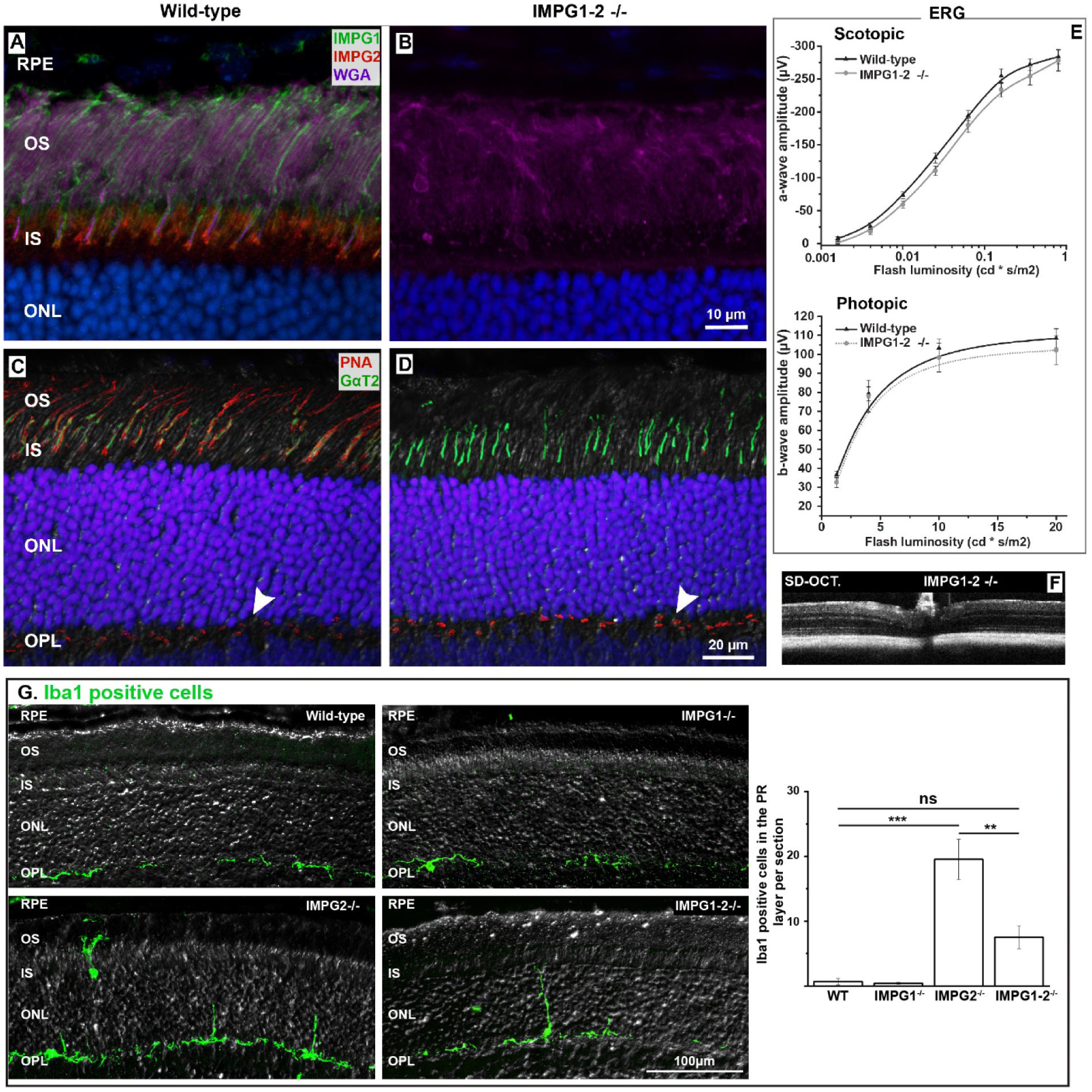
IMPG1-2 double KO mice show an intact retinal structure, function, and lack of subretinal lesions. (A) Wild-type retina stain against IMPG1 (green), IMPG2 (red), and WGA (magenta). (B) IMPG1-2 double KO retina lacks IMPG1 and IMPG2 and shows an unaltered staining pattern for WGA. (C) Wild-type retinas show PNA lectin (red) surrounding the cone OS marker, cone transducin GαT2 (green). (D) Recognition of the cone matrix by PNA is lost in the retina lacking IMPG1-2. Interestingly, staining of the outer plexiform layer (OPL) (arrow) is unaltered. Staining for cone transducin (GαT2) shows normal number of cones in IMPG1-2 double knockouts. Representative images from three eye sections from thee different littermates at P45. (E) Electroretinogram analysis of 8-months-old animals shows no significant differences in scotopic or photopic responses between mutant and wild-type mice. (F) Spectral Domain Optical Coherence Tomography (SD-OCT) representative image of 8-month-old IMPG1-2 double KO mouse showing normal anatomy and the absence of subretinal vitelliform lesions. (G) Evaluation of microglial migration to the photoreceptor layers on wild-type, IMPG1, IMPG2 and double KO mice retinas stained by Iba1 antibody (green) and differential interference contrast (DIC) technique. Retinal sections obtain from five different animals from three different littermates at 5-months-old. Bars are the mean ± SEM (**P <0.01, ***P < 0.001).

Combined, these results indicate that the accumulation of IMPG1 proteoglycan is a key component in the formation of the subretinal lesions. Furthermore, the lesion leads to reactive gliosis and microglia migration to the photoreceptor layers and subretinal space.

### Photoreceptor function is reduced in the absence of IMPG2

To study whether the material accumulated between the OS and RPE and whether the focal subretinal lesion impact on the photoreceptor function, we assessed electroretinograms (ERG) (Fig. 3H, I). The ERG technique measures changes in the retina electric field in response to light flashes stimulation and is capable of differentially test rod or cone response by preadapting the animals to dark (scotopic conditions) or light (photopic condition) respectively. IMPG2 KO mice show a significant decrease in scotopic and photopic ERG responses at different light intensities at 8 months. In contrast to IMPG2 KO mice, IMPG1 KO mice did not show a reduction in the ERG. In 8-month-old animals, the response was similar to wild-type littermate controls.

### Mice lacking IMPG1 and IMPG2 do not develop subretinal lesions or visual deficits

To evaluate the role of IMPG1 in the pathology of IMPG2 null animals and to study possible functional redundancy among IMPGs, we generated IMPG1 and IMPG2 double KO mice. The absence of IMPG1 and 2 proteins in the retina of double KO mice were confirmed by IHC analysis (Fig. 4A, B). We also stained retinas obtained from double KO and wild-type mice with the lectin WGA wich stains jet undescribed IPM components. Interestingly, IMPG1-2 double KO retinas did not show material accumulation, subretinal lesions, or changes in the IPM (Fig 4B). Next, we focused on cones by staining with the intracellular cone OS marker, cone transducin α-subunit, GαT2 (green), and the cone-specific extracellular matrix marker PNA lectin (red) (Fig. 4C, D). There were no changes in the number of cones in the retina lacking IMPGs as judged by transducin expression. However, the PNA staining was absent in the IPM of the IMPG1-2 double KO (Fig. 4D).

We next used photopic and scotopic ERG to analyze the function of the IMPG1-2 double KO photoreceptors (Fig. 4E) and *in vivo* SD-OCT imaging to assess the presence of subretinal lesions (Fig. 4F). Interestingly, these mice did not show any functional deficits or associated subretinal lesions up to 8-months old.

Next, we studied the microglial activation of wild-type, IMPG1 KO, IMPG2 KO, and double KO in retinal sections by counting the Iba1+ cells that migrated to the photoreceptor layers (Fig. 4G). Only IMPG2 KO retinas showed a significant increase of Iba1+ cells.

These results indicate that the cone-specific lectin PNA stains IMPG1 and 2 proteoglycans. In addition, these results show that the absence of both IMPG proteoglycans does not affect the retinal morphology and function. More importantly, removing IMPG1 protein along with IMPG2, strongly reduce the pathological signs observed in IMPG2 KO, suggesting that the mechanism behind visual impairment in IMPG2 KO mice involves the mislocalization and accumulation of IMPG1.

### IMPG1 and 2 are chondroitin sulfate proteoglycans

IMPG proteoglycans bind ChS (Clark et al., 2011, Hollyfield et al., 1999, Lazarus & Hageman, 1992). To confirm those results and to obtain an independent assessment of IMPG mislocalization, we investigated the pattern of ChS localization in the IPM of IMPG mutant animals (Fig. 5). ChS consist of hundreds of repeating disaccharide units including chondroitin 6-sulfated disaccharide (C6S). To expose the C6S epitopes, we treated the retinal tissue with chondroitinase ABC (Caterson, 2012, Caterson, Christner et al., 1985, Couchman, Caterson et al., 1984). Then, we used the anti-stub antibody 3B3 to label the C6S epitopes. The antibody can also stain “native” C6S epitopes in a terminal end of glucuronic acid without chondroitinase ABC treatment. Retinas without chondroitinase ABC treatment show “native” 3B3 epitopes at the outer plexiform layer and sclera but does not show any staining at the IPM (Fig 5 lower panel).

**Figure 5.**
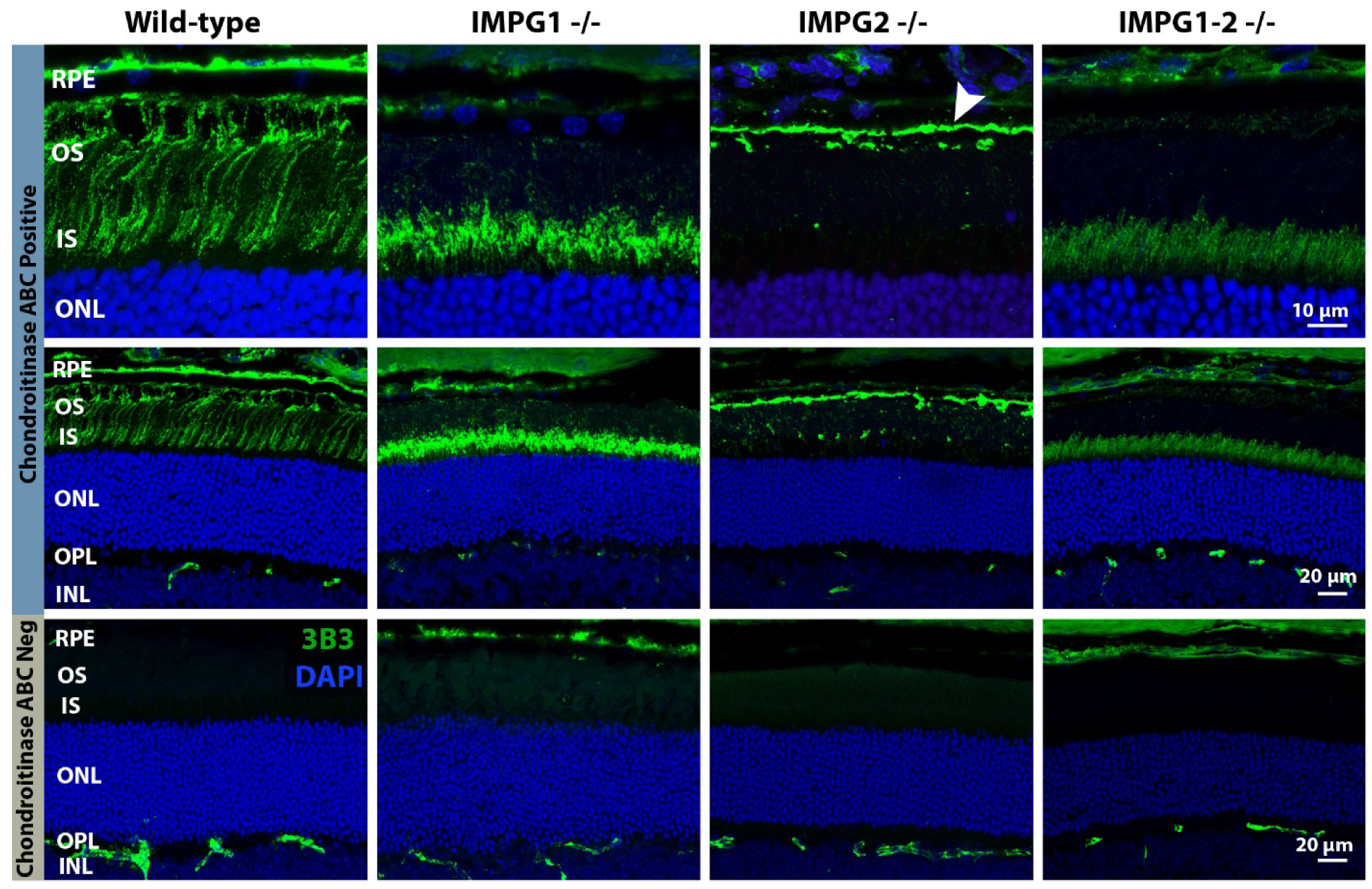
Mislocalization of chondroitin sulfate in the IPM of IMPG1, IMPG2, and IMPG1-2 double KO mice. Immunohistochemistry on retina cross-sections from wild-type and IMPGs KO mice stained with the anti-chondroitin sulfate antibody 3B3 (green), in sections treated with chondroitinase ABC (upper and middle panels) or untreated (bottom panel). Upper and middle panel (100x and 40X magnification respectively): Wild-type retina showing 3B3 staining at the outer section of the IS and through the entire OS, in IMPG1 KO mouse 3B3 staining is restricted to the IS while IMPG2 KO show ChS accumulation at the OS-RPE boundary (arrowhead) plus a scattered staining pattern along the IPM. Both single mutants mimic the mislocalization staining pattern seen for PNA and IMPG1 in figure 3. IMPG1-2 double KO expresses faint staining for 3B3 at the IS. Bottom panel: Retinas untreated with chondroitinase ABC do not show 3B3 staining at the IPM. Images processed from three different animals from three different littermates at P45.

Wild-type retina predigested and stained with 3B3 anti-stub antibody revealed that the IPM of the mouse retinal tissue is rich in ChS and is localized at the IS and OS of the photoreceptors (Fig. 5 upper panels). In the retina lacking IMPG1, ChS is reduced at the OS but shows a robust presence in the IS. On the other hand, in the absence of IMPG2, ChS is mislocalized at the boundary between OS and RPE. In addition, we observed sporadic staining along the OS and at the IS-OS intersection. The observed ChS mislocalization pattern in IMPG mutants are analogous to the PNA and IMPG distribution seen in figure 2. These results suggest that IMPG proteoglycans are covalently modified by ChS and that both IMPGs are responsible for the major staining pattern observed for ChS in the IPM. In addition, the ChS distribution in IMPG2 KO mice points out that the material that accumulates between the OS and RPE is rich in ChS.

Interestingly, in the retina lacking IMPG1 and 2, the ChS staining does not entirely disappear; instead, we found faint staining for ChS at the IS. This finding suggests that other extracellular proteoglycans may compensate for the deficit of ChS in the absence of both IMPG proteins.

## Discussion

In this study, we used knockout technology in mice to determine the contributions of IMPG1 and IMPG2 to the structure and function of the IPM. Our data indicate that both IMPG1 and IMPG2 proteoglycans surround the photoreceptors, wherein IMPG2 facilitates the localization of IMPG1 in the IPM. Moreover, in the absence of IMPG2, IMPG1 mislocalizes generating subretinal lesion comparable to subretinal lesions found in humans and likely causing defects in photoreceptor function.

The proteoglycans IMPG1 and IMPG2 are synthesized by photoreceptors (Chen et al., 2003, Kuehn & Hageman, 1999a, Lee et al., 2000). This finding is supported by our RNA-seq data that show robust expression of IMPG1 and 2 in wild-type retina (Murphy, Cieply et al., 2016). In comparison, in the retina that lack photoreceptor cells (AIPL1^-/-^, Aryl-hydrocarbon-interacting protein-like 1), the expression of IMPG1 and 2 were negligible (IMPG1: WT 1425 vs AIPL1-/- 5; IMPG2: WT 567 vs AIPL1-/- 7, expression data in arbitrary units). We identified distinct compartmentalization of these molecules in the extracellular matrix. IMPG2 remains at the photoreceptor IS distal region anchored to the cell membrane with a putative transmembrane helix. In contrast, IMPG1 was found along the OS region, up to the RPE (Fig. 1 and *SI appendix*, movie 1). In IMPG2 KO retinas, IMPG1 is mislocalized and mostly accumulated in the OS-RPE interface (Fig 2 and *SI appendix*, movie 2). These findings imply that alteration at the IPM around the IS (IS-IPM) provokes deficits in the IPM around the OS (OS-IPM) suggesting that the IS-IPM is involved in the generation of the OS-IPM (Fig. 6).

**Figure 6.**
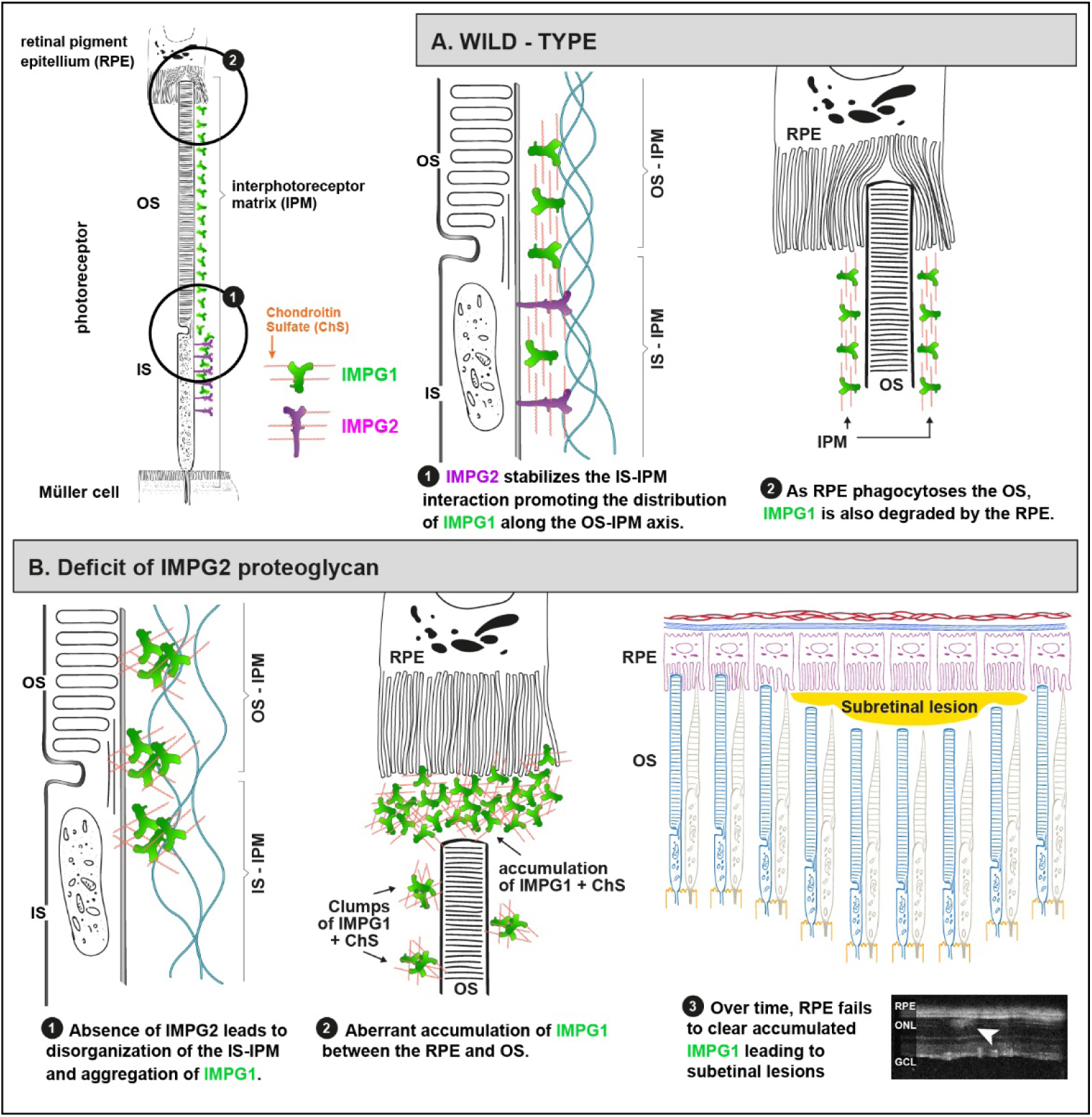
Proposed mechanism of interaction between IMPG1 and IMPG2 in the generation of IPM and the mechanism leading to vitelliform lesions in the absence of IMPG2. A) Wild-type panel show IMPG2 (magenta) anchored to the photoreceptor IS cell membrane facing the IPM and exposing the chondroitin sulfate (ChS). IMPG2 proteoglycans interact with components of the IPM around the IS (IS-IPM) to arrange a specific IS-IPM spatial conformation that allows secreted IMPG1 to align in an ordered conformation. This ordered configuration of IMPG1 allows the protein to follow the OS maturation until it gets phagocytized by the RPE. B) IMPG2 KO panel: Lack of IMPG2 produces altered IS-IPM conformation with no guidance for the assembly of secreted IMPG1. Eventually, unorganized IMPG1 bundle together forming clumps rich in ChS that interfere with the normal IPM clearance by the RPE. This forms a material that accumulates at the subretinal space and activates microglia. Over time, the RPE fails to remove the accumulated material and forms vitelliform lesions (arrow).

Lectins are proteins that bind specific carbohydrates. Two commonly used lectins to identify cone- and rod-photoreceptor-specific matrix are PNA and WGA, respectively (Hollyfield, Rayborn et al., 1990, Hollyfield, Varner et al., 1990, Johnson & Hageman, 1991, Tien et al., 1992). PNA recognizes galactosyl (β-1,3) N-acetyl galactosamine (Galβ1-3GalNAc) residues in glycoproteins (Cummings & Etzler, 2009), while WGA binds to the sialic acid commonly present around neurons (Monsigny, Roche et al., 1980, van der Want, Klooster et al., 1997). Our IHC study showed that the PNA staining colocalizes with both IMPG1 and 2 (Fig. 1) and follows the mislocalized IMPG distribution in IMPG1 and IMPG2 KO mice (Fig. 2). In addition, staining for PNA was abolished in IMPG1-2 double KO (Fig. 4). These findings agree with previous studies that showed that PNA-lectin binds to IMPG1 and IMPG2 in the IPM. (Acharya et al., 2000, Hollyfield et al., 2001). Interestingly, although both rod and cone photoreceptors produce IMPG1 and 2, PNA is used as a cone matrix marker across multiple species. We speculate the reason for lack of IMPG1/2 recognition of rod photoreceptor matrix is due to epitope masking of terminal sugar by sialyl residues. This is supported by studies showing that the removal of sialic acid by treatment eliminated the preferential binding of PNA to cone IPM (Uehara, 1993, Uehara, Ozawa et al., 1995). In contrast, the staining pattern for WGA that recognizes sialic acid was unaltered in our animal models suggesting a robust quantity of sialylated proteins in the IPM.

Chondroitin Sulfate (ChS) is a primary component of numerous ECM biological tissues (Mikami & Kitagawa, 2013). The ChS are linear polysaccharide chains composed of repeats disaccharide units of glucuronic acid and N-acetylgalactosamine. The nomenclature of the ChS is based on the position of sulfur groups substituted on the ChS backbone. There are three different types of ChS sulfation in humans retinas, unsulfated, 4-sulfated and 6-sulfated ChS(Clark et al., 2011). However, in mouse retinas, 6-sulfated and weak 4-sulfated ChS were described (Hollyfield et al., 1999). Our findings show that ChS follows the anomalous IMPG localization pattern in single IMPGs KOs. In addition, we observed a substantial reduction of ChS staining in the IPM of IMPG1-2 double KO (Fig. 5) suggesting that IMPG proteoglycans are the primary source of 6-sulfated ChS at the IPM.

Our results show that IMPG proteoglycans are not essential for the development of functional rods and cones, indicating that these proteoglycans are not key players for OS development and maturation in mice (Fig. 3, 6). Surprisingly, IMPG1-2 double KO mice showed faint ChS staining at the IS, leaving the OS unstained (Fig. 5). We hypothesize that the absence of these normal constituents of the IPM initiates compensatory mechanisms that help stabilize the IPM in the double KO mice. It is likely that the expression of some other ChS proteoglycan is upregulated to compensate for the loss of IMPGs. The exact mechanism behind this phenomenon is currently under investigation.

Humans with missense or nonsense mutations on IMPG2 are linked to two visual disorders, retinitis pigmentosa with the involvement of macula and AFVD. Macular involved retinitis pigmentosa is characterized by macular abnormalities in the context of retinitis pigmentosa (Bandah-Rozenfeld et al., 2010, van Huet et al., 2014). On the other hand, AFVD are characterized by round or oval, yellow, subretinal lesions forming an egg yolk-like vitelliform macular lesion associated with a relatively normal ERG (Alten & Eter, 2015, Brandl et al., 2017, Chowers et al., 2015, Guziewicz et al., 2017, Hamel, 2014, Khan & Al Teneiji, 2019, Meunier et al., 2014, Sivaprasad et al., 2016, Wilde et al., 2016). The reason behind the variability in the symptoms observed in humans is not clear. Our work shows that deficiency in IMPG2 proteoglycan in mouse models produces lesions like those found in humans with IMPG mutations. Humans with mutations in either IMPG proteins develop subretinal lesions in areas with high concentration of cones (Brandl et al., 2017, Manes, Meunier et al., 2013, Meunier et al., 2014). We speculate that cone-specific glycosylation due to IMPG is necessary for the stability of the cone-IPM. In this sense, the lack of macula in our animal model represents a limitation.

Young animals lacking IMPG2 (P45) show early accumulation of material between the OS and RPE that is rich in IMPG1 and ChS (Fig 2, 3, 5), and microglia cells migrating to the photoreceptors layer (Fig 4G). As the animals age, IMPG2 KO mice (5 months) develop subretinal lesions and reduction of the ERG response (8 months) (Fig. 3). These findings along with our observation in animals lacking only IMPG1 and both IMPG1 and 2 (Fig. 2, 4) that do not present protein mislocalization or pathological signs seen in IMPG2 KO mice, suggest that the impairment of IMPG1 to integrate into the IPM and eventual accumulation in the subretinal space is the main cause of subretinal lesion formation and vision loss in IMPG2 KO mice.

Humans with heterozygous missense mutations in IMPG1 also developed vitelliform macular dystrophies (Brandl et al., 2017, Manes et al., 2013, Meunier et al., 2014). Patients with truncated or dysfunctional IMPG1 proteins due to missense mutation of IMPG1 gene present subretinal vitelliform lesions. The findings from this work suggest that the abnormal IMPG1 proteins may not assemble properly in the IS-IPM, mislocalize, and accumulate, similar to our observation in the IMPG2 KO animal model.

Lesions in the central nervous system lead to an increase in the synthesis and deposition of ChS proteoglycans helping to form a glial seal, this phenomenon is also known to occur in retinal degenerative diseases (Dyck & Karimi-Abdolrezaee, 2015, Singh, Kolandaivelu et al., 2014). IMPG proteins are highly glycosylated, the excessive accumulation of IMPG1 covalently bound to ChS at the RPE-OS boundary can, over time, forms something akin to glial seal that, on time, overwhelms the RPE uptake capacity leading to the formation of subretinal lesions and visual impairment. This hypothesis is in line with the fact that patients with defects in the metabolism of GAGs due to abnormal matrix metalloproteinases develop retinopathy characterized by the accumulation of GAGs at the subretinal space (Ashworth, Biswas et al., 2006, Fenzl, Teramoto et al., 2015).

A closer inspection of the subretinal lesions found in our IMPG2 KO mice reveals a dome shape mass that detaches the retina and occupies the subretinal space. These lesions stains with IMPG1, WGA, and present a high concentration of microglia (Fig 3, *SI Appendix* Fig. S3). The retina around the lesions shows signs of damage reactivity and degeneration as microglia migration, reactive gliosis and a reduction of the nuclear cell layer. We also observe that the OS and IS layers close to the lesions present round shape structures that specifically stains with WGA (Fig. 3E.2 and *SI Appendix* Fig. S3). Due to this finding, we also analyzed retina cross-sections in all KO groups. We found that IMPG2 and IMPG1-2 double KO mice show an increased amount of WGA-round shape structures (*SI Appendix* Fig. S4). These structures are similar to ones discovered as a result of photoreceptor degeneration after retinal detachment (Okunuki et al., 2018). There is increasing evidence that relates deficits in extracellular matrix with degenerative disorders like macular degeneration (Al-Ubaidi, Naash et al., 2013, Fernandez-Godino, Pierce et al., 2016). We speculate that IMPGs proteoglycans may play an important role in the photoreceptor normal aging process.

In summary, our findings demonstrate that aberrant localization of IPM proteoglycan is involved in the development of subretinal lesions. Our work proposes that the accumulation of IMPG1 proteoglycan in between the OS and the RPE in IMPG2 KO mice retinas, overloads the RPE clearance capacity and provokes, over time, the failure of RPE, generating the subretinal lesions found in patients (Fig. 6). Overall, our work suggests the existence of a dynamic, interconnected and robust IPM that supports the RPE-photoreceptor microenvironment. The insights provided from this study are essential in deciphering the importance of the matrix in degenerative photoreceptor diseases and in the identification of novel therapies to treat matrix associated blindness.

## Materials and methods

### Animal model

The animal models used in this study were generated by Cripsr-Cas9 technology as described by Moye et. al. (Moye, Singh et al., 2018). Briefly, 20-nt single guide RNA (sgRNA) was synthesized (Clontech Cat.#631441) from 69-nt forward primers against IMPG1 (5’-GCG GCC TCT AAT ACG ACT CAC TAT AGG GGA TCT TTT GGT TCG AAG CTT GTT TTA GAG CTA GAA ATA GCA-3’) and IMPG2 (5’-GCG GCC TCT AAT ACG ACT CAC TAT AGG GGG CGT GAT GAA TAT CGT CAC GTT TTA GAG CTA GAA ATA GCA-3’) synthesized by IDT company (*SI Appendix* Fig. S1). *In Vitro* transcribed sgRNA were used along with Cas9 nuclease (Invitrogen-Cat.#B25641)) and injected into pronuclei of B6D2F1 blastocysts. Heterozygote animals were backcrossed with C57BL/6J (Jackson laboratory). All animals were generated at Transgenic Core of West Virginia University and did not contain *rd1* or *rd8* alleles and are RPE65met homozygous. IMPG1 and IMPG2 KO animals were crossed to produce IMPG1-2 double KO. The animals were maintained under 12-h light / 12-h dark-light cycles with food and water provided *ad libitum*. Both sexes were used in all experiments. All experimental procedures involving animals in this study were approved by the Institutional Animal Care and Use Committee of West Virginia University.

### Immunohistochemistry (IHC)

Mice were euthanized by CO_2_ exposure and eyes were enucleated. A small hole on the cornea was made and the enucleated eyes were placed in 4% paraformaldehyde (PFA) for 30 min. After fixation, cornea and lens were removed and the resulting eye cup was further fixed for 30 min. The eyecups were then washed three times for 5min each in Phosphate-buffered saline (PBS), and left in PBS containing 20% sucrose overnight. The eyecups were placed in 1:1 mix of 20%sucrose and optimal cutting temperature compound (OCT) (Sakura) for two hours, and finally flash-frozen in OCT and stored at −80 °C.

The frozen retinal tissue was sectioned at 16 μm thickness using Leica CM1850 Cryostat, mounted on Superfrost Plus slides (Fisher Scientific) and stored at −20 °C. Retina sections were washed in PBS and incubated for 1 hour in blocking buffer (PBS with 5% Goat Sera, 0.5% TritonX-100, 0.05% Sodium Azide) at room temperature, and then treated with primary antibody overnight at 4° C. On the next day, the sections were washed 3 times for 10 min in PBS-T (0.5% TritonX-100)This was followed by the addition of secondary antibody for two hours at room temperature. The dilutions used for lectins, primary, and secondary antibodies are presented in *Supplementary Information (SI),* Table S1. Sections were washed 3 x 10 min in PBS-T, then sealed with a coverslip after the addition of ProLongGold (Life Technologies).

RPE and retina flat-mount preparations were made from enucleated eyes as described before. Briefly, the cornea was removed after 30 min fixation in 4% PFA and the resulting eyecups were fixed for two more hours. After fixation, eyecup was cut from the periphery towards the optic nerve at four places, after which the retina and RPE were carefully detached under a dissection microscope. The tissue was further processed as described earlier.

Images were acquired and processed with Nikon C2 laser scanning confocal using excitation wavelengths of 405, 488, 543 and 647 nm. Four sections were imaged per sample and data were derived from 3 independent experiments from different littermates.

Enucleated eyes were shipped in fixative for serial cross-section and hematoxylin and eosin staining to Excalibur Pathology Inc. Images were taken with MIF Olympus VS120 Slide Scanner.

The results shown are representative findings that were reproduced in a minimum of three different littermates.

### Photoreceptor ultrastructure

Mice from three different littermates per group were euthanized by CO_2_, and eyes carefully enucleated. Eyes were first fixed in 2% paraformaldehyde, 2.5% glutaraldehyde, 0.1M cacodylate buffer, pH 7.5 for 30 minutes, then cornea and lens were removed and put back in the same fixation buffer at room temperature for 48 hours under rotation (nutator). Following dissection, toluidine blue staining, embedding, and transmission electron microscopy were performed at Robert P. Apkarian Integrated Electron Microscopy Core (IECM) at Emory University, US as described previously (Dilan, Moye et al., 2019).

### Spectral domain optical coherence tomography (SD-OCT)

Animals were anesthetized by isoflurane [2.0% isoflurane at 1 liter per minute oxygen flow rate] and eyes were topically dilated with a 1:1 mixture of tropicamide: phenylephrine hydrochloride. Animals temperature was controlled by a heat pad and the cornea moisture applying ophthalmic gel (Genteal, Alcon). The equipment used for spectral domain optical coherence tomography (SD-OCT), is an Envisu R2200 high-resolution equipment. The images cover an area of 1.4×1.4mm surface centered at the optic nerve with 100 b-scans per eye with 1000 points resolution per b-scan.

### Electroretinogram (ERG)

Adult mice after 24 hours of dark-adaptation mice were anesthetized in an induction chamber with isoflurane 1.5 % at 1 liter/minute oxygen flux. After induction, pupils were dilated with a 1:1 mixture of tropicamide: phenylephrine hydrochloride for 10 minutes prior to recording. The mice were placed on a heated platform and connected to an anesthesia cone mask at the same oxygen flux, the reference electrode placed on the neck and the ground electrode on one leg. Full scotopic flash intensity ERG was performed with white light in the dark-adapted mice, after that, for the photopic ERG mode, a white background light of 30 cd/m^2^ for 10 minutes was used to light adapt animals and four increasing flash intensities were used to stimulate the eyes. ERGs were measured using Celeris system, from Diagnosys LLC. with fully-integrated electrodes built into the stimulator. Eyes were moist regularly to avoid dryness (Genteal, Alcon) and to maintain low impedances.

### Statistics

All data are presented as mean ± standard error margin. ERG responses were analyzed with two-way ANOVA and then Tukey post-hoc test for comparison of means between groups. Iba1 cell count and WGA round shape structures count was analyzed with one-way ANOVA and then Tukey post-hoc test for comparison of means between groups.

## Acknowledgments

The authors thank Thamaraiselvi Saravanan and Angelica Jacques for support in the maintenance of animal models and genotyping, Lucia Bonifacini for aiding in preparation of figures and to Peter Mathers and the transgenic core in generation of the animal models. This work was supported by Knight Templar Eye Foundation (EMS) and National Institutes of Health Grants RO1 EY028035 (VR), R01 EY025536 (VR), R21 EY027707 (VR), West Virginia Lions and Lions Club International Foundation.

## Supporting Information appendix

**Figure S1:**
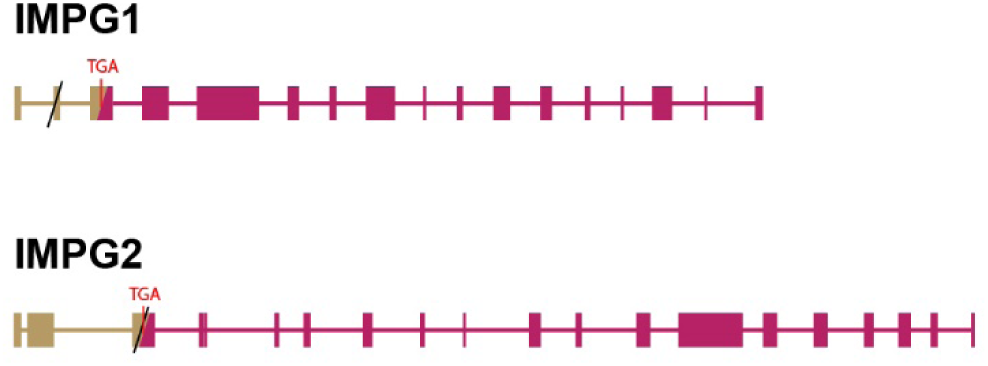
Schematic representation of the change induced in the IMPG1 and IMPG2 knockout mice. Cripsr-Cas9 technology induced deletions are marked by the forward slash (/) symbol and predicted premature termination is represented by TGA codon. Pink colored boxes represent untranslated exons in the mutants beyond the TGA codon.

**Figure S2:**
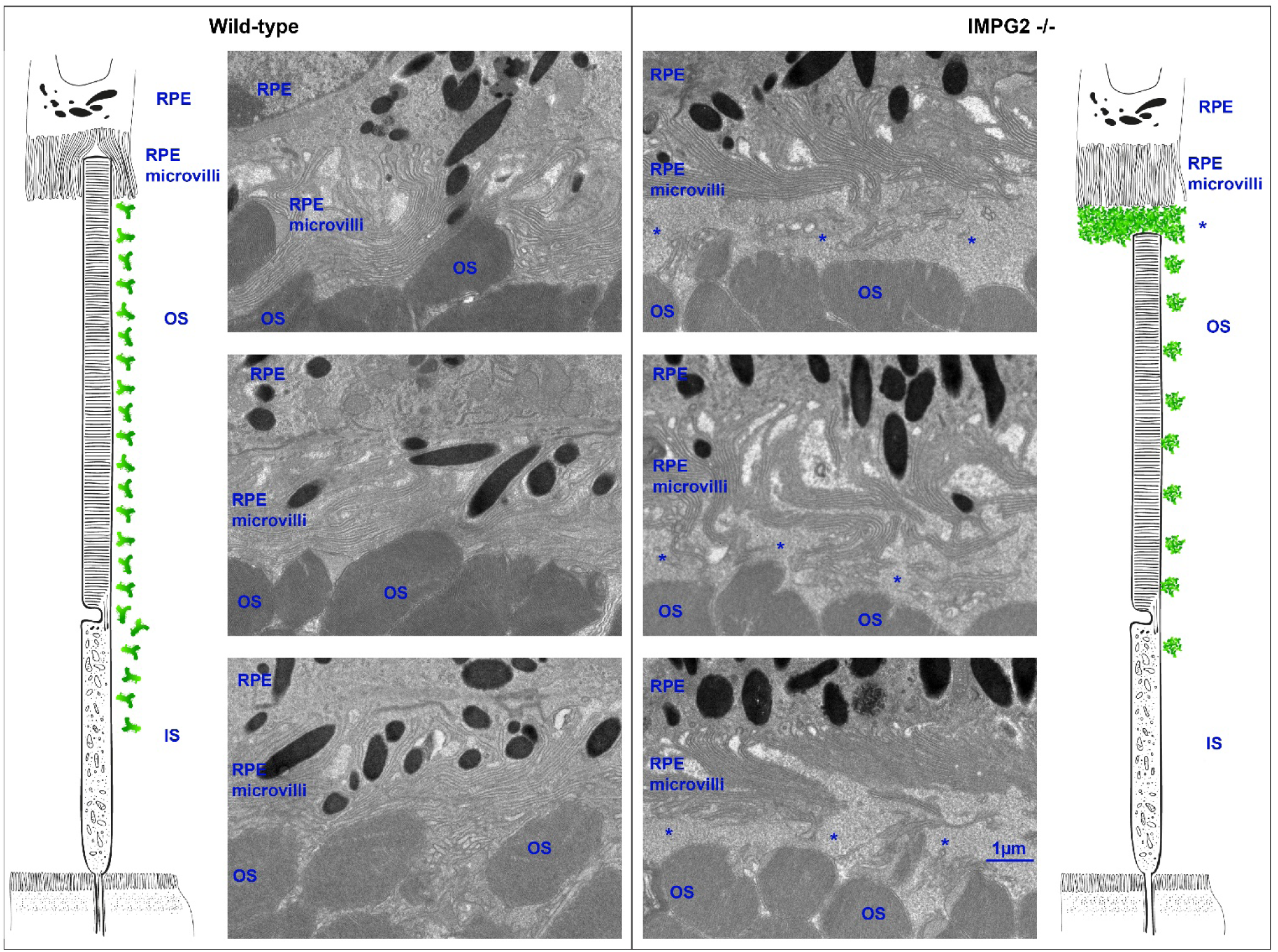
Anomalous material accumulation between photoreceptors outer segments (OS) and RPE in the absence of IMPG2. Electron microscopy images of wild-type retinas exhibit a close contact between OS and microvilli of the RPE. In contrast, IMPG2 KO retinas show a separation between OS and RPE filled by an unstructured material (asterisks). Retinal tissue used was collected from 5-month-old mice (n=3).

**Figure S3.**
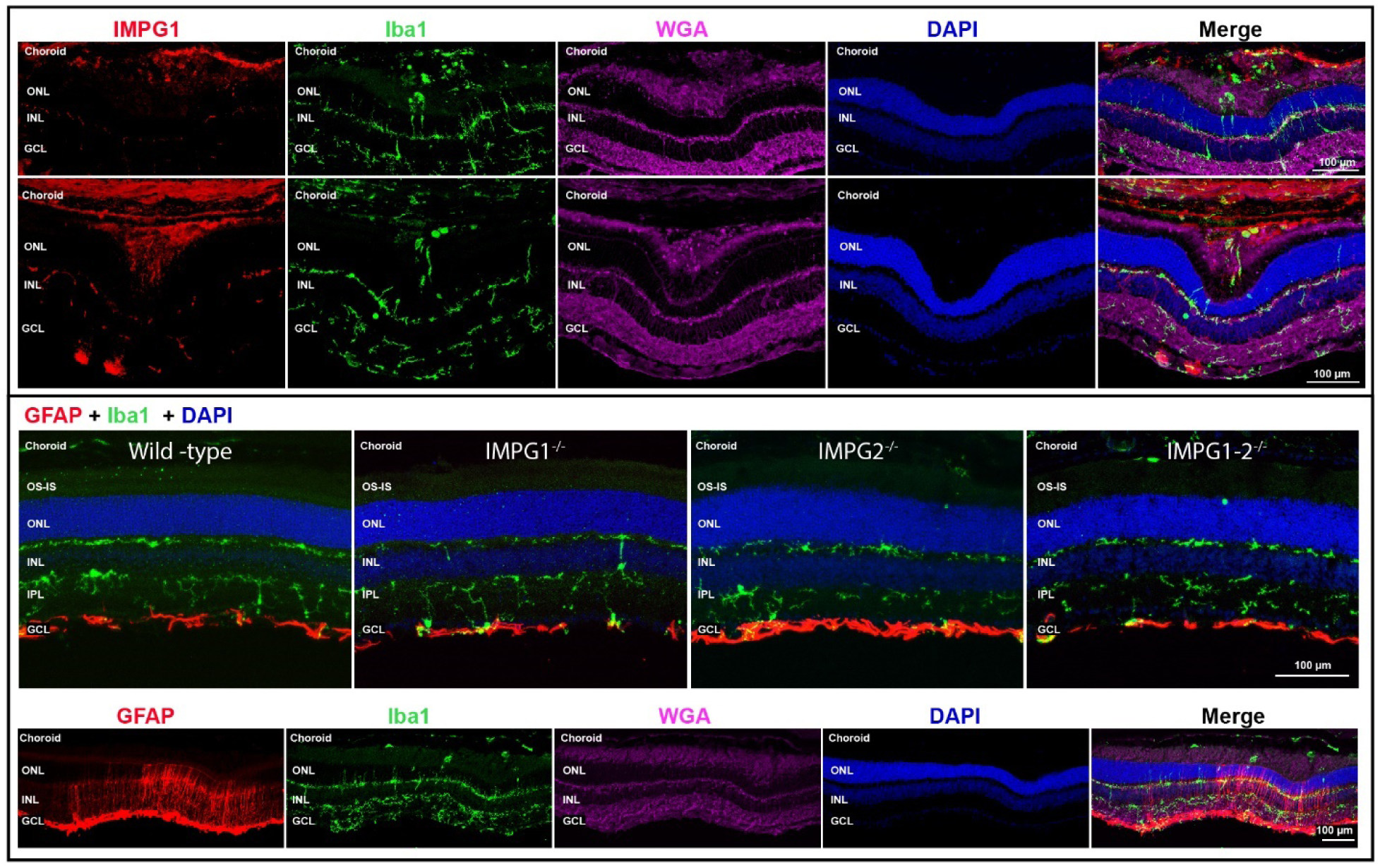
Subretinal lesions show IMPG1 accumulation, microglia migration, and glial reactivity. Top panel: Retina cross-sections from two IMPG2 KO animals with subretinal lesions showing IMPG1 (red) and microglia/macrophage stained with Iba1 antibody (green) in the subretinal space. Lower panel: The top figure shows normal glial fibrillary acidic protein (GFAP) (red) staining astrocytes in the ganglion cell layer (GCL) from wild-type, IMPG1 KO, IMPG2 KO, and double IMPG1-2 KO mice retinas. The bottom figure shows a low magnification subretinal lesion in IMPG2 mice showing Müller glia reactivity expressing GFAP limited to the area of the subretinal lesion.

**Figure S4.**
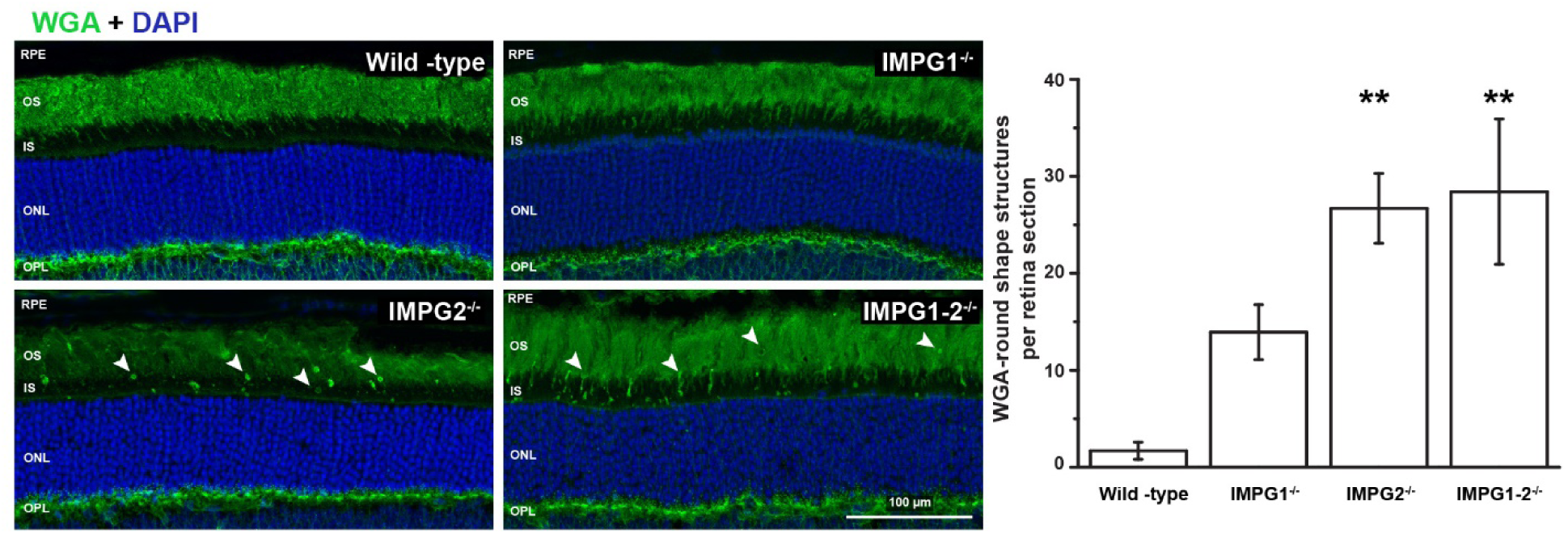
WGA lectin positive puncta at the photoreceptors IS and OS of IMPG2 and IMPG1-2 double KO mice. Left panel, representative retina cross-section of Wild-type, IMPG1, IMPG2, and IMPG1-2 double KO mice stained with WGA (green) lectin. Distinctive puncta (arrows) used for quantification on the left side. Images processed from five different animals from three different littermates at 5-months-old. Bars are the mean ± SEM (**P <0.01).

**Table S1.** List of names and specifications of antibodies and lectins used in this work.

| Name | Host | Against | Dilution | Source | Catalog |
| --- | --- | --- | --- | --- | --- |
| SPARC (G-11) | mouse | IMPG1 / SPARC | 1:2000 | Santacruz | SC-377366 |
| Anti-IMPG2 | rabbit | IMPG2 / SPARCAN | 1:1000 | Sigma | HPA008779 |
| ZO-1 | rabbit | ZO-1 | 1:500 | Invitrogen | 61-7300 |
| GαT2 (I-20) | rabbit | cone transducine | 1:1000 | Santacruz | SC-390 |
| Iba1 | rabbit | iba1 protein | 1:1000 | Wako | 019-19741 |
| 3B3 | mouse | Chondroitin 6 sulfate | 1:100 | Amsbio | 270433-CS |
| GFAP-Cy3 | rabbit | GFAP | 1:1000 | Sigma | C-9205 |
| PNA |  | N-acetylgalactosamine | 1:1000 | Vector lab | B-1075 |
| WGA-680 |  | N-acetyl-D-glucosamine | 1:1000 | Invitrogen | W32465 |
| WGA-488 |  | N-acetyl-D-glucosamine | 1:1000 | Invitrogen | W11261 |

**Movie S1 (separate file).** Wildtype retina flat mount 3D reconstruction centered in one cone, stained with PNA (red), IMPG1 (green), IMPG2 (magenta), DAPI (blue). PNA colocalizes with IMPG2 around the IS and with IMPG1 around the OS. IMPG1 stains weakly at the IS in association with IMPG2 proteoglycan.

**Movie S2 (separate file).** 3D reconstruction of the flat-mount retina from wild-type, IMPG1, and IMPG2 KO mice, stained with PNA (red), IMPG1 (green), Gαt2 (magenta), DAPI (blue). Wildtype shows Gαt2 in the center of the cone photoreceptors surrounded by PNA and IMPG1. In IMPG1 KO retina, PNA localizes just in the base of the cone, and Gαt2 antibody stains the cone OS. In IMPG2 KO retina Gαt2 stain the cones OS surrounded by scattered staining pattern for IMPG1.

